# DBDA matrix increases ion abundance of fatty acids and sulfatides in MALDI-TOF and mass spectrometry imaging studies

**DOI:** 10.1101/2023.01.13.524013

**Authors:** Nigina Khamidova, Melissa R. Pergande, Koralege C. Pathmasiri, Rida Khan, Justin T. Mohr, Stephanie M. Cologna

## Abstract

MALDI-TOF MS is a powerful tool to analyze biomolecules owing to its soft ionization nature and generally results in simple spectra of singly charged ions. Moreover, implementation of the technology in imaging mode provides a means to spatially map analytes in situ. Recently, a new matrix, DBDA (N1,N4-dibenzylidenebenzene-1,4-diamine) was reported to facilitate the ionization of free fatty acids in the negative ion mode. Building on this finding, we sought to implement DBDA for MALDI mass spectrometry imaging studies in brain tissue and successfully map oleic acid, palmitic acid, stearic acid, docosahexaenoic acid and arachidonic acid using mouse brain sections. Moreover, we hypothesized that DBDA would provide superior ionization for sulfatides, a class of sulfolipids, with multiple biological functions. Herein we also demonstrate that DBDA is ideal for MALDI mass spectrometry imaging of fatty acids and sulfatides in brain tissue sections. Additionally, we show enhanced ionization of sulfatides using DBDA compared to three different traditionally used MALDI matrices. Together these results provide new opportunities for studies to measure sulfatides by MALDI-TOF MS including in imaging modes.

## Introduction

Matrix assisted laser desorption/ionization-time of flight mass spectrometry (MALDI-TOF MS) has enabled the rapid analysis of biomolecules from an array of sample types owing to the soft ionization nature and generation of primarily singly charged ions. Moreover, with the advent of imaging strategies, MALDI mass spectrometry imaging (MSI) has enabled spatial mapping of biomolecules, drugs and other analytes in situ. One such class of biomolecules, lipids, which are critical in cellular structure integrity, signaling, and energy storage have been analyzed broadly using MALDI MSI strategies. For example, fresh frozen bone samples have been imaged resulting in discovery of unique signatures associated with bone marrow and soft tissues^1^, and phospholipids have been mapped in a *Drosophila* model of amyotrophic lateral sclerosis brain tissue^2^. Other examples include lipid mapping in cancer models^3^, ischemic stroke^4^, and of the human retina ^5^.

Despite some common physiochemical properties of lipids, generally they are a class of biomolecules that are quite diverse. Fatty acids serves as the blocks of other lipids however, free fatty acids are endogenously generated by lipolysis and have functional cellular roles. MSI has been performed on different model systems to investigate the spatial distribution of free fatty acids. For example, dynamics of isotopically labeled DHA and AA were monitored in brain tissue following administration in mice ^6^. Moreover, fatty acids and their respective double bond positions have been mapped in cancer tissues ^7^.

Another family of bioactive lipids is sulfolipids and one class includes sulfatides (SHexCer) which are sulfated glysosphingolipids. The biological functions of sulfatides is vast and in the brain, sulfatides are highly enriched in myelin ^8^. With regard to MSI, sulfatides have been mapped in brain tissues to understand oligodendrocyte development ^9^ as well as in renal tissues of a model of Alport Syndrome ^10^. Thus, continued mapping of these crucial lipids is of utmost important in biological specimens.

While MSI has been a successful strategy in measuring lipid distributions, the heterogeneous nature of lipids requires utilization of differing matrixes and ionization models. Common MALDI matrices for lipid analysis include: 1,5-Diaminonaphthalene (DAN), α-Cyano-4-hydroxycinnamic acid (CHCA), 2,5-dihydroxybenzoic acid (DHB), 9-aminoacridine (9-AA). In a recent study, the compound N1,N4-dibenzylidenebenzene-1,4-diamine (DBDA), was shown to ionize small molecules (<500 Da) including free fatty acids without matrix interference ^11^. In this study, we investigate the usage of DBDA to acquire MALDI-MS images of free fatty acids from murine mouse brains, as well as sulfatides from mouse brains. Finally, we compare the ion abundance of sulfatide standards with commonly used MALDI matrices versus DBDA.

## Materials and methods

### Reagents

Sulfatides standards were obtained from Avanti Polar Lipids (Alabaster, AL). LC-MS grade Acetonitrile (ACN), Methanol (MeOH), 9-Aminoacridine (9-AA), α-Cyano-4-hydroxycinnamic acid (CHCA), peptide standards bradykinin 2-9, angiotensin I and II, ACTCH 18-39 and ammonium formate were obtained from Sigma-Aldrich (St. Louis, MO). 1,5-Diaminonaphthalene (DAN) was obtained from TCI Chemicals (Portland, OR). Tissue-Tek O.C.T. compound was received from Sakura Finetek (Torrance, CA). N1,N4-dibenzylidene benzene-1,4-diamine (DBDA) was synthesized as previously described ^11^. *Experimental model:* All experiments were performed in accordance with the University of Illinois Chicago IACUC approved protocols. Heterozygous Balb/c *Npc*^*nih*^ (*Npc1*^*+/ −*^) mice were obtained from Jackson Laboratories (RRID: IMSR JAX:003092) and a breeding colony was maintained in our laboratory. Genotyping was performed using polymerase chain reaction as previously described ^12-13^. Control (*Npc1*^*+/+*^) mice at ten weeks of age were euthanized by CO_2_ asphyxiation and subsequently decapitation. Whole brain tissue from control and mutant mice were dissected and immediately frozen in dry ice to maintain spatial integrity and stored at −80°C until further use.

### Sample preparation

A CryoStar NX50 Cryostat (Thermo Fisher Scientific) was used to cryosection frozen, intact brain tissue at −11°C. Eight, serial sagittal sections from wild type and mutant brain were obtained at 10-μm thickness and thaw-mounted directly on the same MALDI stainless steel target plate, then stored at −80 °C until further analysis. Tissues were thawed and washed in cold 50 mM ammonium formate solution for 15 seconds. The tissues were vacuum dried at room temperature and weighed prior to matrix application.

### Matrix Sublimation

*Matrix Sublimation*: A homemade sublimation apparatus was used to apply MALDI matrix to produce an even coating of N1,N4-dibenzylidene benzene-1,4-diamine (DBDA). Forty milligrams of DBDA were solubilized in 3 mL of acetone, then aspirated onto the bottom of sublimation flask. Acetone was evaporated using a N_2_ stream to form an even layer of matrix on the bottom of the sublimation flask. The hotplate was set to 128 °C, and to monitor the actual temperature, a digital thermometer probe was placed in contact with the bottom of the sublimation flask. An ice slush was placed in the cold finger which the MALDI plate was adhered on the underside using copper tape. Sublimation was done under 80 mTorr pressure for 12 minutes. Matrix was applied at a density of 28 mg/cm^2^.

### Mass spectrometry dried-droplet analysis

Sulfatide standard mix was dissolved in methanol. DAN, 9-AA, and CHCA were dissolved in 70% ACN/H_2_O, and DBDA was dissolved in 80% ACN/H_2_O resulting in 5 mg/mL of matrix concentration. A 1:1 (v/v) ratio of matrix and a solution of 0.01 nmol/μL sulfatide standard were mixed before spotting on the MALDI plate. MALDI-TOF/TOF mass spectrometry was performed using a Sciex 4800 MALDI-TOF/TOF mass spectrometer equipped with 200 Hz Nd-YAG (355 nm) laser and was externally calibrated with a peptide calibration mix in the positive and negative reflectron mode. Six replicate measurements were taken for 9 different sulfatide species. Lipid data was acquired in negative ion reflectron mode 750-1000 *m/z* range. MALDI-MS spectra were collected on dried droplet mixture of 5 mg/mL DBDA and 1nmol/μL ST mixture. Seven replicates were analyzed for 21 different ST mixture dilutions using 3000 laser intensity.

### MALDI mass spectrometry imaging

Data acquisition was carried out as previously reported. In brief, the raster size was set to 100 µm using 4800 Imaging Tool v3.2 (https://ms-imaging.org/wp/4000-series-imaging) and laser shots per a pixel was set to 50. Data was acquired in negative ion reflection mode with the mass range 750 – 1000 m/z for sulfatides and 200-500 m/z for free fatty acids. Data processing was done using MSiReader v1.0 ^14-15^. GraphPad Prism 9 was used for statistical analysis.

## Results and Discussion

Imaging lipids to visualize their spatial distribution is not widely feasible using antibody-based strategies and mass spectrometry imaging has been a powerful tool to enable such endeavors. A challenge with lipid analysis is the need for optimized matrix consideration for different classes of lipids. As previously reported, the highly stable, DBDA matrix is an excellent choice to ionize fatty acids in a dried-drop experiment^11^. In the current study, we aimed to further analyze the utilization of DBDA as a matrix for lipid MSI and to explore the possibility of enhanced ion abundance of another lipid class namely, sulfatides. The experimental workflow is depicted in **Figure S1**.

### Spatial distribution of free fatty acids using DBDA as a matrix

We first sought to determine the appropriateness of carrying out MALDI-MSI using DBDA as the matrix for free fatty acids mapping in whole brain tissue. Using sagittal murine brain sections, we demonstrate that indeed we observe strong signals for free fatty acids (**Figure 1A)**. Notably, we observe clear isotopic distributions of known fatty acid masses, which are labeled in the order of palmitic acid (16:0) m/z 255.2, oleic acid (18:1) m/z 281.2, stearic acid (18:1) m/z 283.3, arachidonic acid (20:4) m/z 303.2, docosahexaenoic acid (22:6) m/z 327.2. All observed ions are [M-H]^-^ and were confirmed with on tissue MS/MS. A representative image with labeled brain features is provided in **Figure 1B**. Next, we generated the MALDI image for free fatty acids in 10-week-old mice (**Figure 1C**). We observe that the majority of fatty acids seem to be localized in the grey matter of the brain. The highest ion signal for all five fatty acids arises from the cerebral cortex and the grey matter of the cerebellum. Palmitic acid (16:0) (PA(16:0)) and stearic acid (18:0) (SA(18:0)) exhibit high ion intensity on the cerebral cortex as well as on the outer regions of the cerebral cortex. Oleic acid (18:1) (OA(18:1)) seems to be in higher abundance in the white matter than the other common fatty acids. Strong signal for docosahexaenoic acid (DHA) is observed to be solely localized on the grey matter of the brain. Based on these data, we can conclude that DBDA can be used to acquire a spatial distribution of fatty acids directly from the tissue with high reliability.

**Figure 1.**
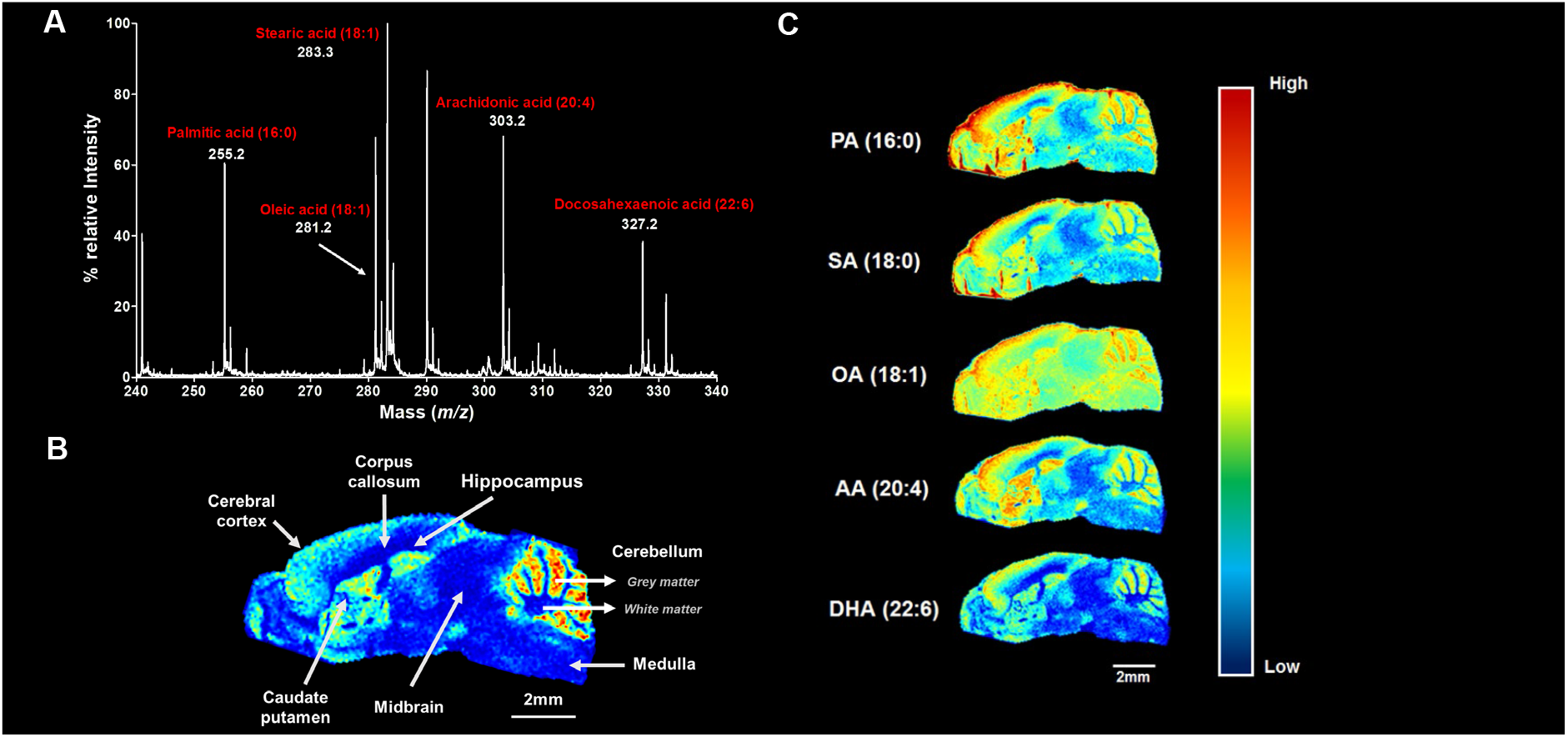
DBDA as a matrix for the MALDI analysis of fatty acids in murine brain tissues. (A) N1,N4-dibenzylidenebenzene-1,4-diamine (DBDA) was applied via sublimation at 128°C for 12 minutes to 10μM brain tissue sections. Shown is the on-tissue MALDI-TOF mass spectrum for *m/z* 240-340 where peaks corresponding to common brain fatty acids are annotated. (B) Ion density map of DHA (*m/z* 327.2) showing that distinct brain regions (C) MALDI-mass spectrometry imaging (MSI) analysis was performed on sagittal cryosections from 10-week mice. Shown are Ion density maps for palmitic (PA), stearic (SA), oleic (OA), DHA and arachidonic (AA) acid.

### Acquiring spatial distributions of sulfatides using DBDA matrix2mmLow

To further analyze the usage of DBDA as a matrix to ionize other lipids, we evaluated the possibility of another anionic lipid class, sulfatides. Firstly, we investigated the on-tissue mass spectrum by acquiring data from m/z 800-910 (**Figure 2A**). Strong signals corresponding to the accurate mass of sulfatide lipids were observed, again as [M-H]^-^ species. We observed nine different sulfatide species with differing abundance. The most abundant sulfatide detected in mice brain tissue was SHexCer(d18:1/24:1) with *m/z* of 888.5 [M-H]^-^ ion. The least abundant sulfatide detected in mice brain in SHexCer (d18:1/20:0) with *m/z* of 834.5 [M-H]^-^ ion. The abundance of the two monoisotopic peaks abundance. Other sulfatides that were observed include SHexCer (d18:1/18:0) with *m/z* of 806.5, SHexCer (d18:1/18:0(2OH)) with *m/z* of 822.5, SHexCer (d18:1/20:0) with *m/z* of 850.5, SHexCer (d18:1/22:0) with *m/z* of 862.5, SHexCer (d18:1/22:0(2OH)) with *m/z* of 878.5, SHexCer (d18:1/24:1(2OH)) with *m/z* of 904.5, and SHexCer (d18:1/24:0(2OH)) with *m/z* of 906.5. The mass spectrum indicated the capability of DBDA to ionize sulfatides thus warranting exploration of imaging strategies. Next, we performed on tissue MS/MS for these precursor ions to confirm the molecular structure and identity. ON tissue MS/MS of *m/z* 878.5 (**Figure 2B**) confirmed the assignment. Key fragment ions observed include HSO_4_- (*m/z* 97.1), sulfated sugar group (SHex, *m/z* 241.1) and 22:0-OH fatty acyl chain (*m/z* 568.4). **Figure 2C** is a representative image for the spatial distribution of the hydroxylated sulfatide SHexCer d18:1/22:0 (2OH). These mass spectra and mass spectrometry images of sulfatides indicate that DBDA can be used as a matrix in MALDI-MSI to obtain spatial distribution of sulfatides.

**Figure 2.**
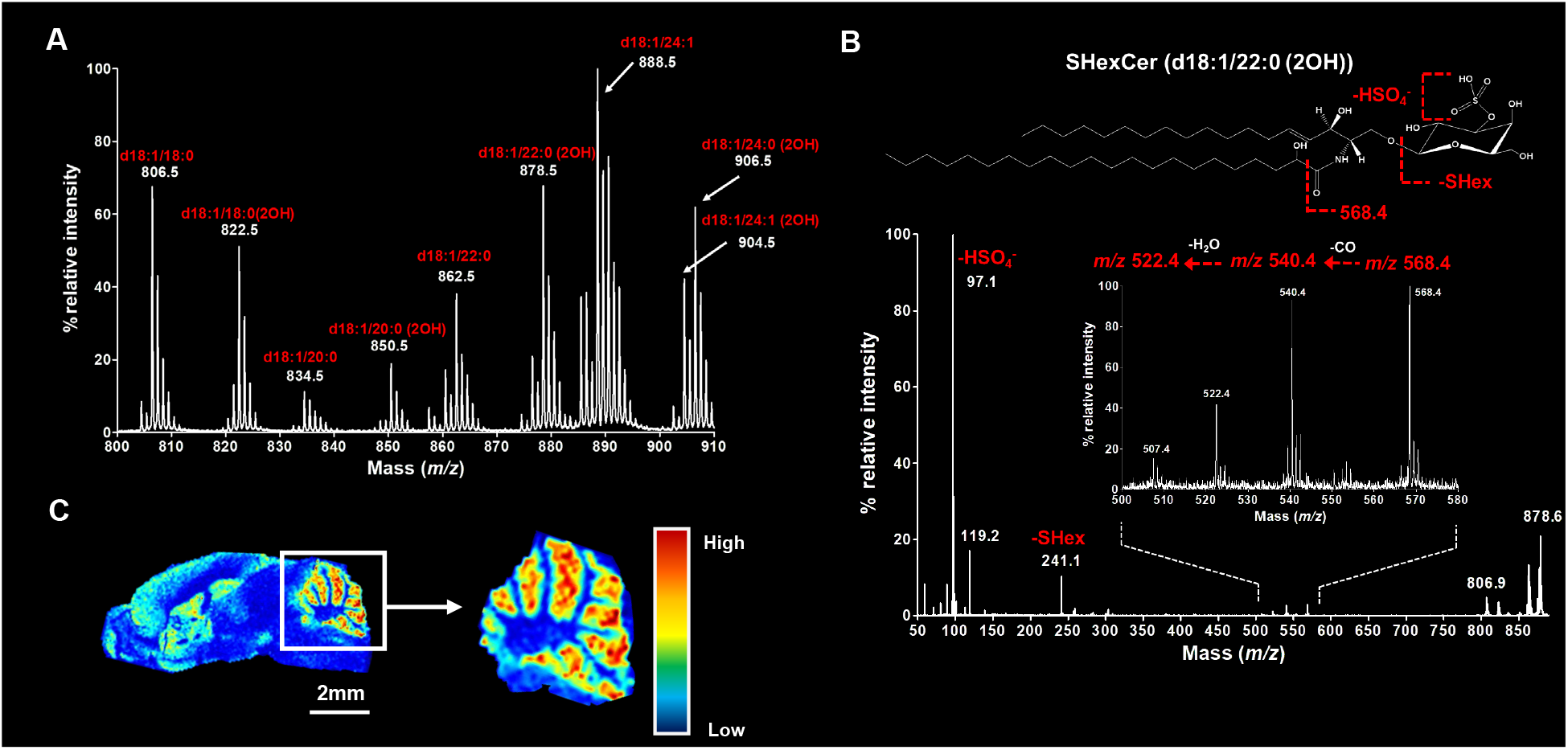
DBDA as a matrix for the MALDI analysis of sulfatides in murine brain tissues. (A) N1, N4-dibenzylidenebenzene-1,4-diamine (DBDA) was applied via sublimation. Shown is the on-tissue MALDI mass spectrum for *m/z* 800-910. Putative mass assignments for multiple sulfatides are annotated. (B) On-tissue tandem MALDI-MS/MS mass spectrum for *m/z* 878.5 where peaks corresponding to fragments are annotated. Here, we observed the loss of the HSO4- (*m/z* 97.1), the sulfated sugar group (SHex, *m/z* 241.1) and 22:0-OH fatty acyl chain (*m/z* 568.4). (C) MALDI-mass spectrometry imaging (MSI) analysis was performed on sagittal cryosections from 10-week mice. Shown is the ion density map for d18:1/22:0 (2OH).

### DBDA enhances ionization of sulfatides compared to other matrices

Numerous studies using differing MALDI matrices to ionize biomolecules have been conducted. As previously reported, DAN facilitates both negative and positive ion production and has been successfully implemented for imaging^16^. Other matrices that are useful for lipid imaging but are challenging to sublime include CHCA and 9-aminoacridine. Despite these challenges, considering the proper MALDI matrix is important to generate strong signals for the ions of interest. Given our positive result for imaging sulfatides with DBDA, we sought to determine if ion abundance was increased compared to other standard MALDI-MS matrices. Using the dried droplet method, a standard mixture of nine of the common sulfatides was analyzed (**Figure 3**). The sulfatide mixture contained SHexCer(18:1/22:0), SHexCer (18:1/20:0)(2OH), SHexCer (18:1/20:0), SHexCer (18:1/18:0)(2OH), SHexCer (18:1/18:0), SHexCer (18:1/24:1),

**Figure 3.**
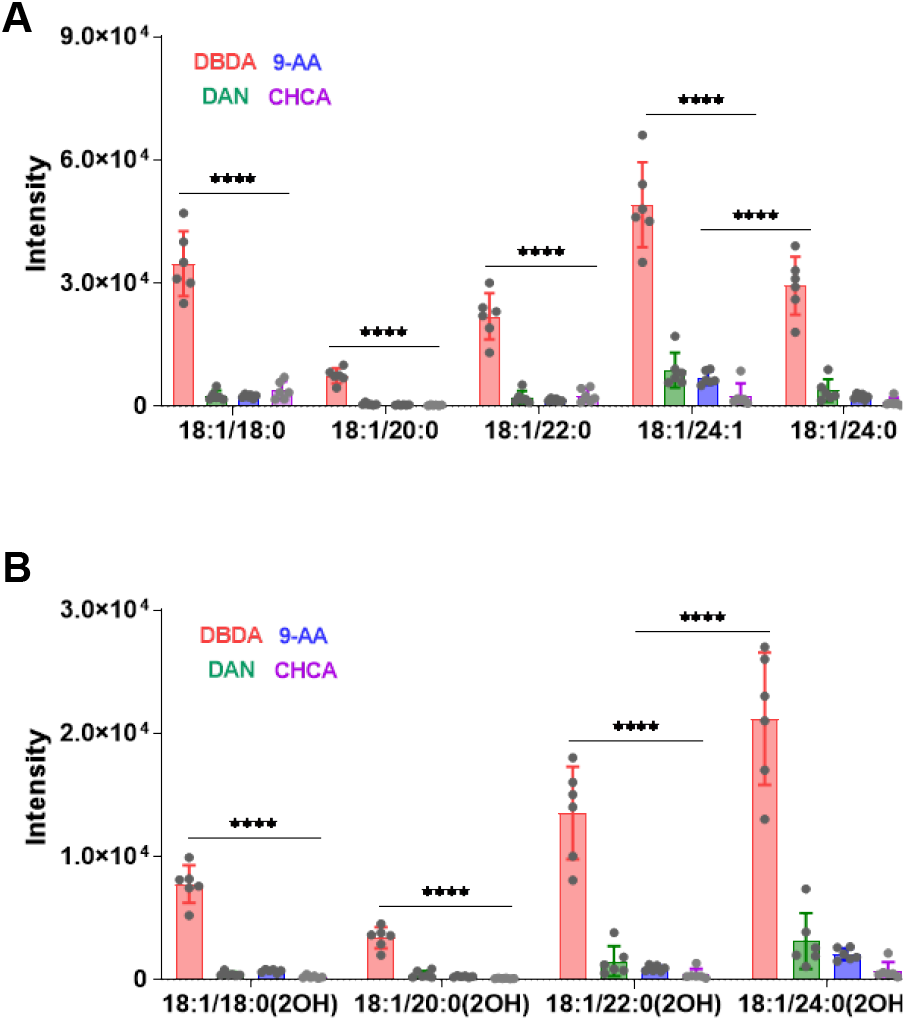
Comparison of different MALDI matrices for the analysis of sulfatides. Standards for both (A) non-hydroxylated and (B) 2-hydroxy sulfatides (B) were analyzed by MALDI-MS using different matricesFor all standards, the use of the DBDA matrix resulted in a significant (one-way ANOVA, p<0.0001 (****)) increased intensity compared to other matrices.

SHexCer (18:1/24:0), SHexCer (18:1/24:0)(2OH), SHexCer (18:1/22:0)(2OH). Examples of MS/MS spectra from this mixture for four lipids is provided in **Figure S2**. The four matrices we sought to compare were: DBDA, CHCA, 9-AA, and DAN. The same solution concentration was used for all analyses. **Figure 3A**, displays a comparison of these matrices for four non-hydroxylated sulfatides. We observe that the non-hydroxylated sulfatides that were analyzed with DBDA matrix exhibit higher ion abundance than other three matrices with high significance, p<0.0001 (****). The same results are seen in hydroxylated sulfatides with 18, 20, 22 and 24 acyl chains (**Figure 3B**) With these data, we conclude that DBDA enhances ionization when compared to DAN, 9-AA, and CHCA.

## Conclusions

In the current study, we sought to utilize DBDA as a negative ion MALDI matrix to image free fatty acids in brain tissues. Moreover, we explore the possibility of performing MSI using DBDA for the anionic sulfated lipids, sulfatides. Our data confirms the feasibility of each.

Finally, we compared the ion abundance of sulfatide standards using DBDA versus other commonly used MALDI matrices and find that the most abundance signal for these standards is observed when DBDA is employed. These data support the notion that DBDA is an excellent MALDI matrix for both free fatty acids and sulfatides using both standard dried droplet methodologies as well as MALDI-MSI via sublimation.

## Acknowledgements

This work was supported in part by the Department of Chemistry, University of Illinois Chicago. Additional funding is acknowledged from the National Institutes of Health, UIC Portal to Biomedical Research Careers (UIC PBRC) PREP (R25GM121212), NIA/NINDS R01 R01NS114413 and the National Science Foundation CAREER Award # 2143920.

