## Supplemental Information for "DBDA matrix increases ion abundance of fatty acids and sulfatides in MALDI-TOF and mass spectrometry imaging studies"

**Address reprint requests to:**

Stephanie M. Cologna, PhD

Associate Professor

Department of Chemistry, University of Illinois Chicago

845 W Taylor Street, Room 4500

Chicago, IL 60607

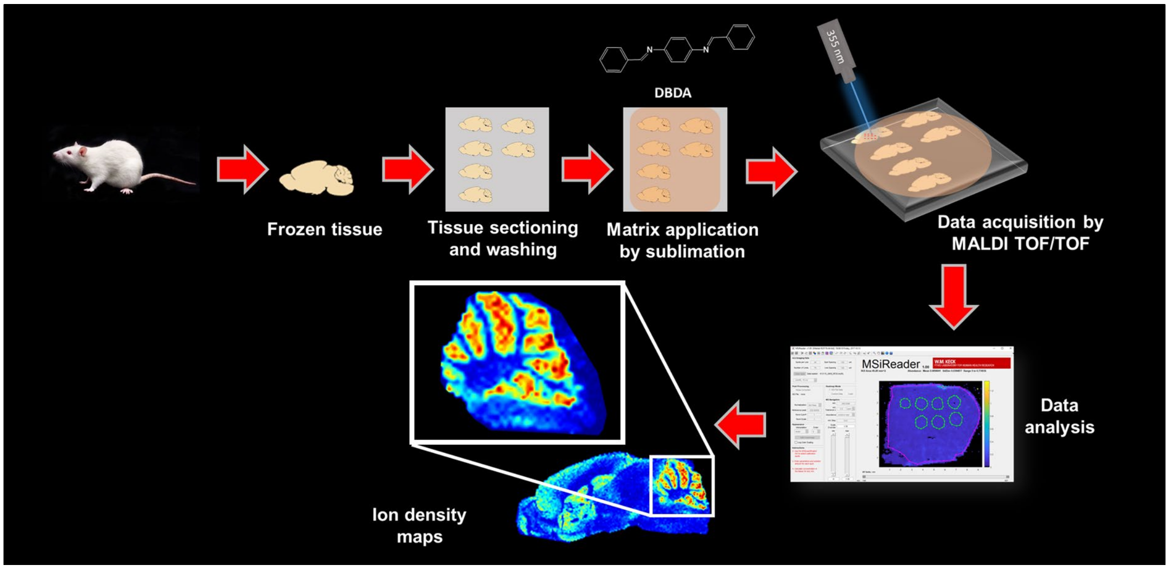


**Figure S1.** Experimental workflow for imaging analysis with DBDA.

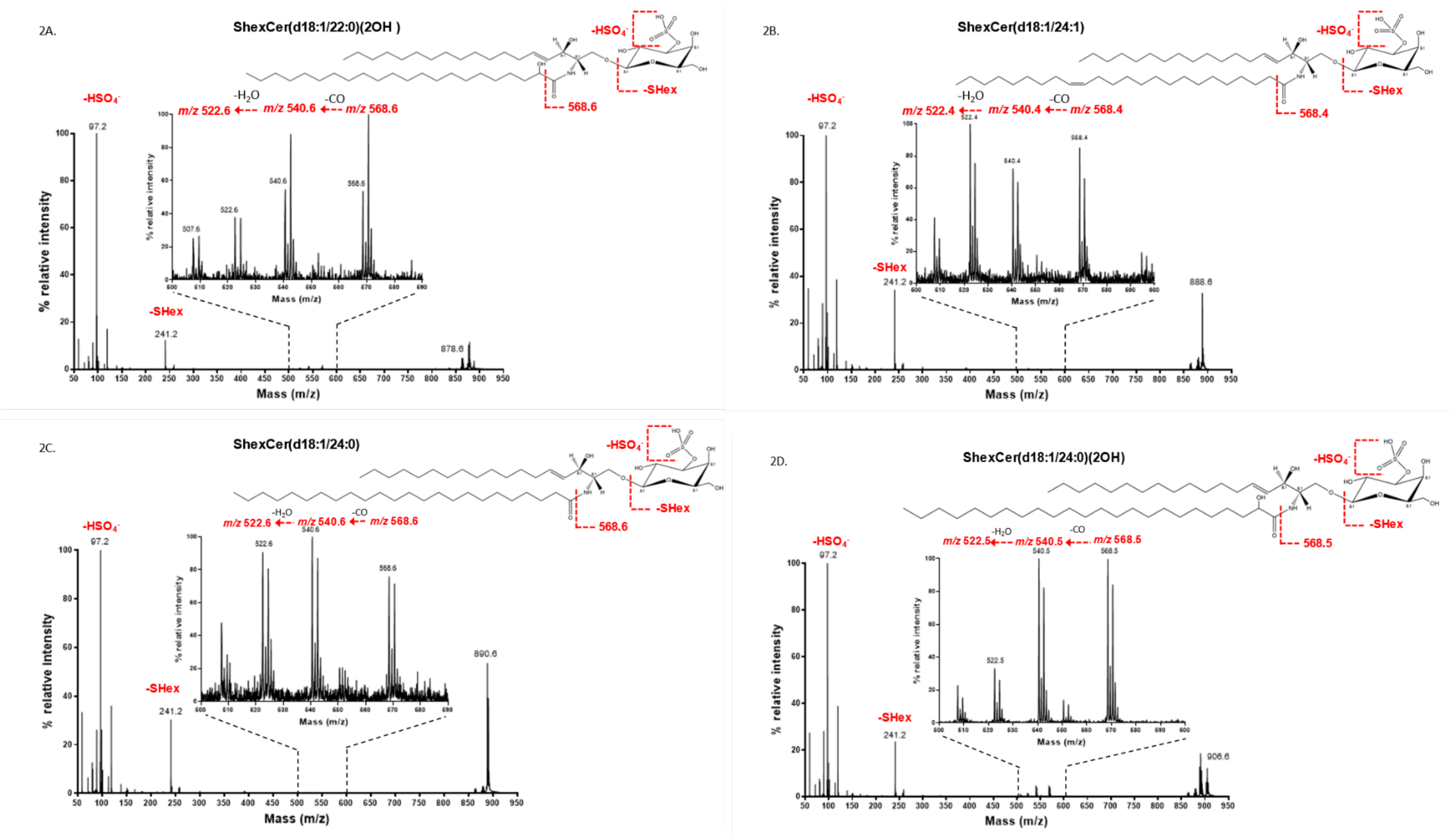


**Figure S2. Tandem MS of sulfatides from standard mixture**. A) Shown is the MS2 with mass spectrum for *m/z* 878.6 where peaks corresponding to fragments are annotated. Here, we observed the loss of the $HSO_4$- (*m/z* 97.1), the sulfated sugar group (SHex, *m/z* 241.1) and 22:0-OH fatty acyl chain (*m/z* 568.4). B) Shown is the MS2 with mass spectrum for *m/z* 888.6 where peaks corresponding to fragments are annotated. Here, we observed the loss of the $HSO_4$- (*m/z* 97.1), the sulfated sugar group (SHex, *m/z* 241.1) and 22:0-OH fatty acyl chain (*m/z* 568.4). C) Shown is the MS2 with mass spectrum for *m/z* 891.6 where peaks corresponding to fragments are annotated. Here, we observed the loss of the $HSO_4$- (*m/z* 97.1), the sulfated sugar group (SHex, *m/z* 241.1) and 22:0-OH fatty acyl chain (*m/z* 568.4). Shown is the MS2 with mass spectrum for *m/z* 906.6 where peaks corresponding to fragments are annotated. Here, we observed the loss of the $HSO_4^-$ (*m/z* 97.1), the sulfated sugar group (SHex, *m/z* 241.1) and 22:0-OH fatty acyl chain (*m/z* 568.4).
